# EXPANSION OF ANTIGEN- SPECIFIC MEMORY CELLS AS A POTENTIAL BOOSTER FOR FOOD TOLERANCE INDUCTION

**DOI:** 10.1101/2023.01.30.526302

**Authors:** Airton Pereira e Silva, Bárbara Oliveira Marmello, Ana Letícia Bentes, Claudia Regina Josetti das Neves Faccini, Sónia Kristy Pinto Melo Rodrigues, João Ricardo Almeida Soares, Sylvia Maria Nicolau Campos, Gerlinde Agate Platais Brasil Teixeira

## Abstract

Approximately 3% of children in Western countries are diagnosed with peanut allergy, a likely lifelong disease. The preferred treatment for food allergy is allergen avoidance. However, oral immunotherapy is an FDA-approved treatment to re-induce tolerance, still, not all patients respond as expected. Thus, the aim of this work is to evaluate whether the association of an antigen-specific tolerogenic (oral tolerance) bystander effect can ameliorate the recovery of inflamed intestinal mucosa. Adult male C57BL/6 mice were divided into five groups, four of which were submitted to an intestinal inflammation induction protocol to peanuts. After sensitization, experimental groups were orally challenged with either peanuts or a hybrid diet (peanuts + mouse chow). In a second stage, groups were sensitized, challenged with peanuts, and then received either peanuts, hybrid diet, or ovalbumin chow during the recovery period of the inflamed mucosa. Results showed no changes in diet intake and body weight. Polyisotypic anti-peanut IgG and IgG1 were significantly increased in the serum from animals in allergic groups. The group that received the hybrid diet showed an increase in CD4^+^CD25^+^Foxp3^+^ regulatory T cells, as well as in B220^+^CD3^-^CD27^+^ memory B cells. Histology of the duodenum showed a decrease in intraepithelial leukocytes in animals who received hybrid diet. Together, our results show that when the tolerogen is added to a diet containing the allergen, it can ameliorate the induction of local inflammation. Simultaneously offering the allergen with a tolerated food increased the mucosal recovery due to the expansion of previously induced memory cells.

## 1. Introduction

Food allergy, which is defined as an immunological adverse reaction to food components, has a high social impact [1]. Understandably, the main route of exposure to food allergens is the gastrointestinal tract, however, antigens may also gain the organism through the skin or the respiratory tract [2]. In allergic individuals, children or adults, clinical manifestations may include dysphagia, food refusal, early satiety, regurgitation, nausea, vomiting, diarrhea, or constipation, etc [3, 4].

Food avoidance is still considered the most effective way for food allergy treatment [5]. However, and especially for children, one may have trouble avoiding unexpected contacts to hidden allergens, [6], i.e., potential allergens not detailed in the ingredients list. One critical potential hidden allergen is peanut [7], with an annual report rate of unintentional ingestion as high as 12% in children and adolescents [8, 9].

Peanut allergy affects 1-3% of children in Western countries and approximately 6% of the US pediatric population [10, 11], with most allergic children remaining allergic for the rest of their lives [12]. Several authors have shown through the years that, although it is possible to induce oral tolerance in children allergic to peanuts, the success rates are relatively low, 18-30% by the end of the treatment [13-16]. The most frequent treatment to induce desensitization in allergic individuals is the oral immunotherapy, which involves the ingestion of progressively increasing doses of the allergen until reaching a target dose (also called eliciting dose) that will then be ingested regularly [17]. The goal is to change the immune response from an allergenic Th2 type to a more tolerogenic Th1 type [16, 18]. Nevertheless, this treatment can trigger anaphylactic reactions, therefore requiring a medical facility, at least during the early stages of therapy [19]. A novel treatment, which was recently approved by the US Food and Drug Administration (FDA) under the commercial name Palforzia [20], has an efficacy of 67%, which is much greater than previously reported treatments in the literature. These participants are those that pass the exit-food-challenge after completing the active treatment. [21] Those patients that do not respond to the treatment, upon cessation are susceptible to an increased chance of resensitization to peanuts [22]. Therefore, it is imperative to better understand the immunological mechanisms involved in oral tolerance induction in previously sensitized patients. Thus, the aim of this work is to evaluate whether the memory cells that induce antigen-specific oral tolerance can interfere collaborating in the kinetics of recovery of the inflamed intestinal mucosa in a mouse model of peanut allergy.

## 2. Material and Methods

### 2.1. Ethics Statement

This work was approved by the Institutional Animal Care and Use Committee under the permit number 00147-09 and follows the ARRIVE guidelines [23].

### 2.2. Animals

Adult C57Bl/6 (60 days old – n=6 animals/group) male mice bred at the Animal Facility of the Federal Fluminense University (Niteroi, RJ, Brazil) were used. Animals were kept in polypropylene cages with stainless steel covers, along with free access to food and HCl-acidified water (pH 2.5) to avoid bacterial contamination and diseases [24, 25]. A conventional environment was maintained with temperature of 25±2.0°C, 60±5% humidity and 12h light/12h dark cycle. To ensure their health, animals were monitored daily for one month prior to the experiment. Body weight was assessed weekly, along with visual inspection of disease symptoms, like labored respiration or shakiness [26, 27].

### 2.3. Induction of the antigen-specific inflammatory gut reaction

Commercial mouse chow (Nuvilab CR1 - NUVILAB-NUVITAL^®^, Sao Paulo, Brazil) containing exclusively corn proteins was offered conforming to the animal facility routine and was used as the tolerogenic food. Animals also received *ad libitum* challenge diet (CD) for 60 days either composed of peanut *in natura* or in-house ovalbumin (OVA) chow according to AIN-93 (report of the American Institute of Nutrition that establishes standards for nutritional studies with experimental rodents [28]). Negative control (NC) group was kept with mouse chow throughout the experiment [29-31]. Food intake was measured three times a week and the mean caloric intake was calculated per gram of body weight (gbw) in each group as previously described [32].

### Antigenic proteins and Immunization protocols

Animals were immunized twice, with subcutaneous injections of 100μg of in house prepared peanut protein extract (PPE) [33], plus 1mg of alum adjuvant [Al(OH)_3_] in the primary immunization and without alum in the booster 21 days later. Animals were bled 200μl from the retroorbital plexus 14 days after each immunization and at the end of CD. Sera were not pooled and were stored at -20°C.

### 2.4. Determination of Antibodies levels

In-house serial dilution enzyme-linked immunosorbent assays (ELISA) were performed to detect specific polyisotypic IgG and IgG1 as previously described [32]. Briefly, after coating 96-well microplates (low binding, Corning, Acton, USA) with 4*μ*g of PPE, a 3-fold serial dilution of each serum (starting at 1/100 v/v) was performed. Goat HRP-anti-mouse IgG or IgG1 (Sigma-Aldrich, Darmstadt, Germany) was used as determined by the manufacturer and reactivity was revealed with OPD (Sigma-Aldrich, Darmstadt, Germany) following specifications and read at 492nm on an automated ELISA reader (Anthos 2010 by Biochrom, London, UK). The relative sensitivity estimate was 95.4% (95% confidence interval = 90.1% to 99.8%).

### 2.5. Determination of T and B lymphocyte profile

Mesenteric lymph nodes (MLN) and spleen (SPL) of each animal were collected, and individually analyzed. In summary: cells were retrieved by mechanical disruption of MLN or SPL onto a sterile steel sieve over a Petri dish containing 5mL of PBS on ice and centrifuged at 300×g for 10 min at 4°C as previously described [34]. Debris were removed, and cell pellets were re-suspended using 3mL PBS on ice. Red blood cells were lysed using hypotonic shock and then removed from SPL samples. After another centrifugation cycle, cell pellets were re-suspended and normalized at 2.0×10^7^ cells/mL using iced PBS. Non-specific binding sites were blocked using 1% fetal bovine serum (Sigma-Aldrich, Darmstadt, Germany) for 30 minutes. Cells were stained with eBioscience™ PerCP anti-mouse CD8 (catalog no. MCD0831 – clone 5H10) along with eBioscience™ Mouse Regulatory T Cell Staining Kit (catalog no. 88-8118-40): FITC anti-mouse CD4 (catalog no. 11-0040 – clone RM4-5), PE anti-mouse CD25 (catalog no. 12-0390 – clone PC61.5) and APC anti-mouse Foxp3 (catalog no. 77-5775-40 – clone FJK-16s). We also used APC anti-mouse CD27 (Biolegend^®^ catalog no. 124211 – clone LG.3A10), FITC anti-mouse CD3 (Biolegend^®^ catalog no. 100305 – clone 145-2C11) and PerCP anti-mouse B220 (Biolegend^®^ catalog no. 103235 – clone RA3-6B2) according to the manufacturer’s dilution instructions. Samples stained with anti-mouse Foxp3 were permeabilized and fixed with buffers from the kit (permeabilization buffer eBioscience™ catalog no. 00-8333-56 and fixation buffer eBioscience™ catalog no. 00-5123-43). After running the samples on BD^®^-C6 Flow Cytometer (Franklin Lakes, USA), all analyses were performed in the lymphocyte gate in FSC (Forward scattering) and SSC (side scattering) plot. We evaluated the following lymphocytes populations: T cells - CD4^+^, CD8^+^, CD4^+^CD25^+^Foxp3^+^ and CD8^+^CD25^+^Foxp3^+^ as well as B cells - B220^+^CD3^-^CD27^+^. Results were normalized to ensure the same cell count per plot (total of 2.0×10^4^ cells per plot) and are shown as percentage of cells in specific gate. Gating strategies are specified in supplemental figure 1.

### 2.6. Histomorphometry of Intestinal segments

All animals received an overdose of anesthetics (60 mg/kg of Xylazine + 350 mg/kg of Ketamine, produced by Sespo Industries^®^, Paulinia, São Paulo, Brazil), after which a longitudinal section was performed to expose the peritoneum. Possible macroscopic alterations were examined in the abdominal cavity. Intestinal segments were collected from each animal and were immediately fixed with 10% buffered formaldehyde and were later stained with hematoxylin-eosin (HE). Tissue sections were scanned using Aperio^®^ Scanscope (Leica Microsystems GmbH, Wetzlar, Germany) and then analyzed with ImageScope software (version 11.2.0.780; Leica Microsystems GmbH, Wetzlar, Germany) to quantify histological parameters. A histomorphometric classification system for normal and inflamed mouse duodenum developed by our group [35] was used to better stage the intestinal situation of each animal.

### 2.7. Experimental groups and timeline

Mice were first divided into three groups. A schematic description of each group is depicted in Table 1:

**Table 1.**
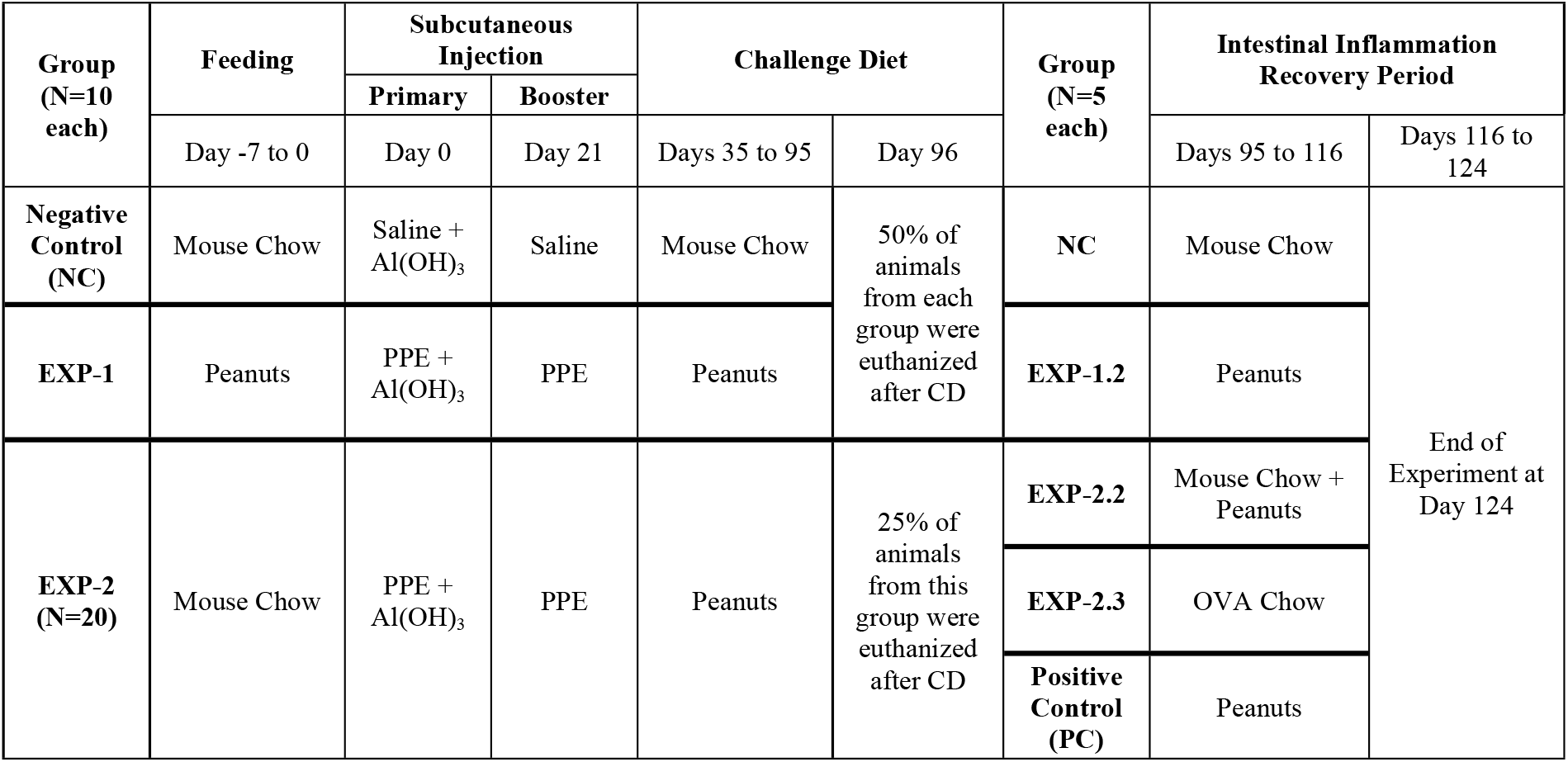
Timeline of experimental protocol.

NC (Negative Control): Mice were weaned and maintained on mouse chow during all periods including the seven days prior to physiological saline inoculations and CD, when animals received mouse chow.

EXP-1 (Experimental 1): Mice weaned on mouse chow received exclusively peanuts for seven days prior to subcutaneous PPE inoculation. On the day of primary inoculation peanuts were removed and switched to mouse chow until the CD, when they received peanuts for 60 days. EXP-2 (Experimental 2): Mice were weaned and maintained on mouse chow during all periods including the seven-day period prior to the inoculation with PPE interval and CD, when they received peanuts for 60 days.

After CD, some of the animals from each group were euthanized for intestinal samples removal. The remaining animals of each group were submitted to an intestinal inflammation recovery period protocol for 21 days. At the end of the treatment period, in addition to the collection of intestinal segments, SPL and MLN, blood was collected from all animals.

NC: Animals kept receiving mouse chow.

PC (Positive Control): Animals received only peanuts in their diet.

EXP-1.2 (Experimental 1.2): A hybrid diet (mouse chow and peanuts) was offered.

EXP-2.2 (Experimental 2.2): Animals were offered an experimental OVA chow as previously described [32]. OVA chow was used to understand the effect of an unrelated protein used both to induce tolerance or allergy during the treatment period.

EXP-2.3 (Experimental 2.3): This group continued receiving peanuts.

### 2.8. Statistical Analysis

Mean ± standard deviation (sd) was used to display the results. First, Shapiro-Wilk test was used for normal distribution assessment. Then, Grubbs’ test was used to remove outliers. Finally, the minimum significance level with 95% confidence interval (p) was calculated using one-way analysis of variance (ANOVA) with Tukey-Kramer post-test (multiplicity adjusted *p*-value). All statistical tests were performed using IBM^®^ SPSS^®^ Statistics version 28.0.1.1 (IBM Corp., Armonk, USA) software. Graphics were built using GraphPad Prism^®^ software version 7.04 (GraphPad Software, Inc., La Jolla, USA) with the same data.

## 3. Results

### 3.1. Results from the First Stage Analyzed

#### 3.1.1. High anti-peanuts polyisotypic IgG and IgG1 titers correlate to allergy induction

Considering all analysis points (post feeding, primary and booster inoculation, and CD) the intergroup analysis revealed that EXP-2 (3.91±1.10) had significantly higher mean polyisotypic anti-peanut IgG levels (p=0.001) when compared to EXP-1 (1.95±0.28) and NC (0.86±0.13), which were not statistically different from each other (p=0.09) (Figure 1A). The same pattern was also observed for anti-peanut IgG1 titers. EXP-2 (1.58±0.76) had significantly higher mean IgG1 levels (p=0.001) when compared to EXP-1 (0.67±1.28) and NC (0.38±0.53), which were not statistically different from each other (p=0.09) (Figure 1B).

**Figure 1.**
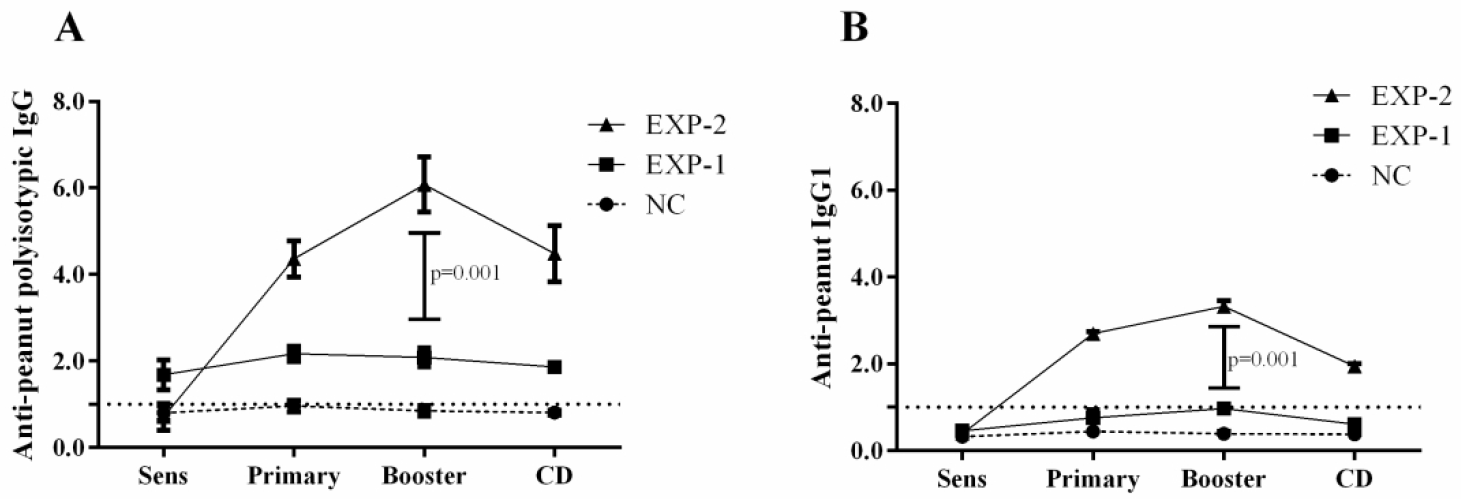
**A)** Mean group polyisotypic IgG. EXP-2 showed significant higher titers when compared to EXP-1 and NC, which was immunized with saline. **B)** Mean group IgG1. EXP-2 showed significant higher titers when compared to EXP-1 and NC, which was immunized with saline. Results were expressed as ELISA index (EI) where values of EI *>*1.0 (horizontal dashed line) were considered positive No pooled sera were used, each individual mouse was analyzed.

#### 3.1.2. Effector and regulatory immune cells in MLN and SPL

After gating the lymphocyte region in MLN samples, we observed that EXP-2 (45.40±3.98) had significantly higher CD4^+^ cell titers in plot (p<0.001) than EXP-1 (33.92±4.25) and NC (36.40±5.43), which were not statistically different from each other (p=0.07) (Figure 2A). In SPL, no significant differences (p=0.80) were observed among NC (20.33±1.15), EXP-1 (19.66±1.13) and EXP-2 (20.66±1.53) (Figure 2A). Representative plots of each group can be found on supplemental Figure 2.

**Figure 2.**
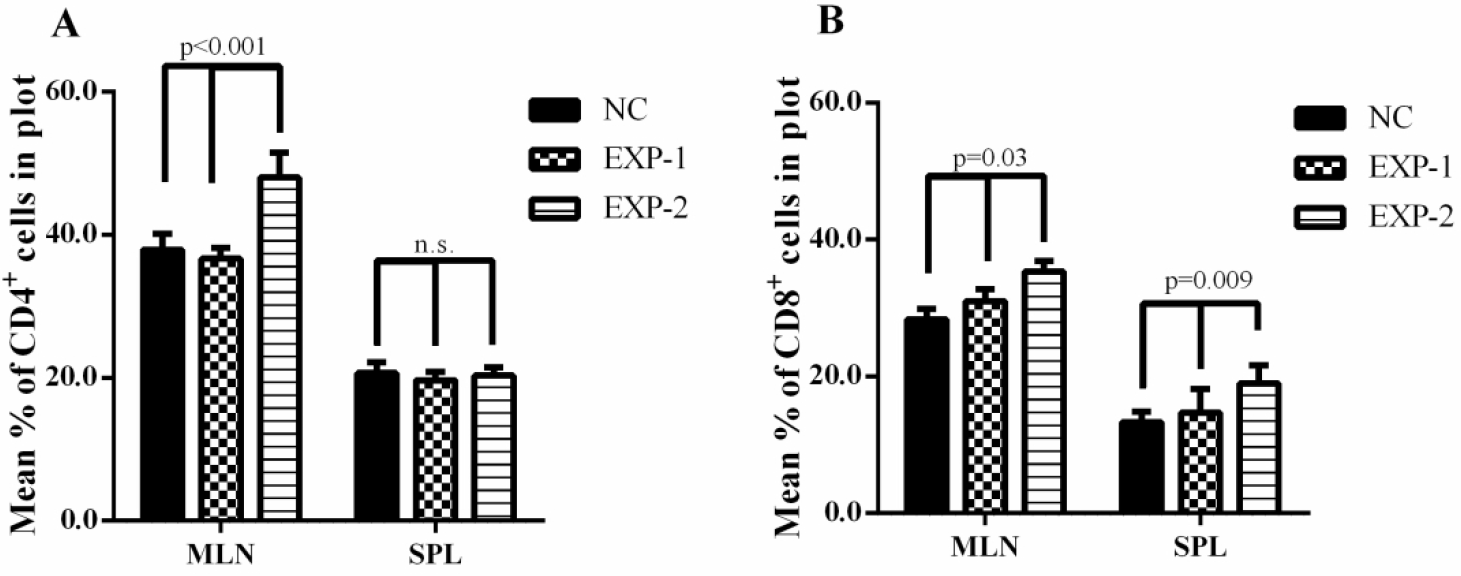
**A**) Mean percentage of CD4^+^ T lymphocytes in MLN and SPL in gate P3 ± SD per group. **B**) Mean percentage of CD8^+^ T lymphocytes in MLN and SPL in gate P3 ± SD per group.

Statistically significant differences were observed among CD8^+^ T cells in MLN, where EXP-2 (35.31±2.67) showed significantly higher titers (p=0.03) than EXP-1 (31.07±1.12) and NC (29.30±1.71) (Figure 2B). Similarly, in SPL, EXP-2 (19.33±1.72) showed significantly higher titers (p=0.009) than NC (13.33±1.52) and EXP-1 (14.33±2.01) (Figure 2B).

The regulatory CD4^+^CD25^+^Foxp3^+^ T cell population analysis in MLN showed that NC (10.36±2.77) presented significantly lower titers (p=0.001) than EXP-1 (17.00±3.45) and EXP-2 (14.67±2.21). EXP-1 also showed significantly higher titers (p=0.02) than EXP-2. Likewise, in SPL, NC (7.23±5.45) showed significantly lower titers (p=0.001) than EXP-1 (12.60±3.59) and EXP-2 (14.06±4.11) (Figure 3A). Regarding CD8^+^CD25^+^Foxp3^+^ T cells, however, NC (0.60±0.10) and EXP-2 (1.53±0.99) showed significantly lower (p=0.002) titers when compared to EXP-1 (2.90±0.25). In SPL, NC (0.56±0.12) showed significantly lower titers (p<0.001) than both EXP-1 (1.41±0.72) and EXP-2 (1.13±0.31). (Figure 3B).

**Figure 3.**
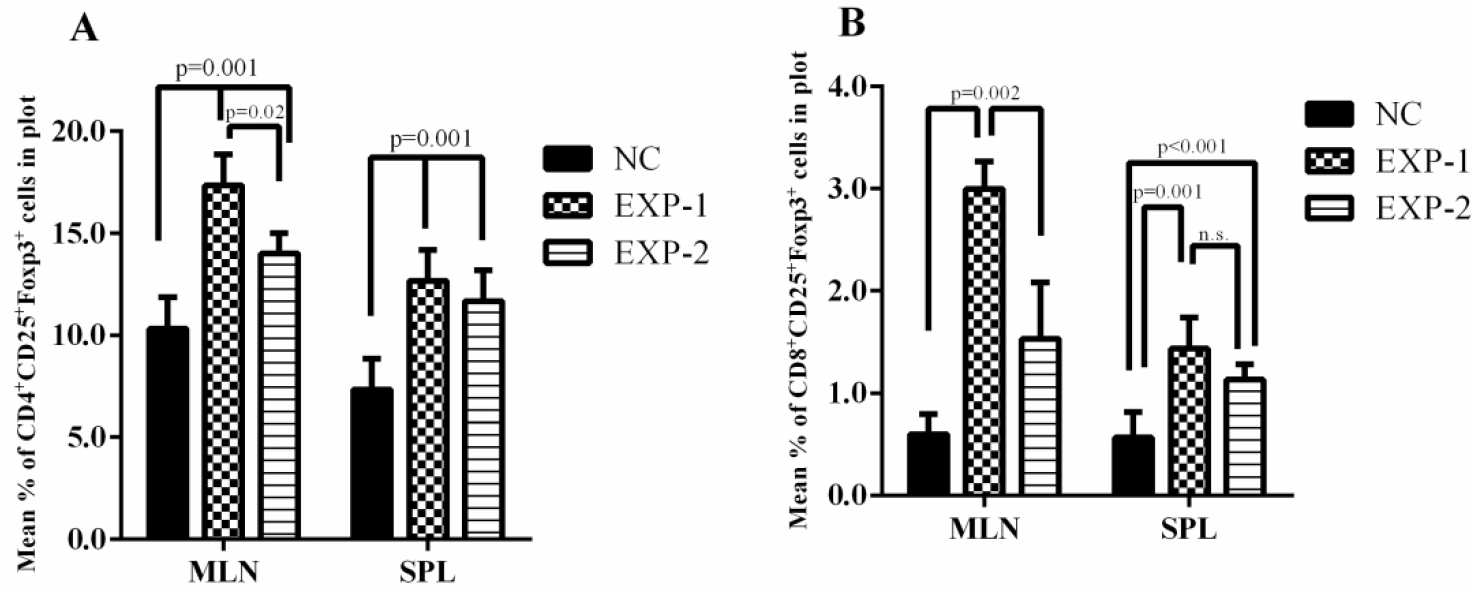
Mean group percentage in gate P3 ± SD of: **A)** CD4^+^CD25^+^Foxp3^+^ T lymphocytes in MLN and SPL. **B)** CD8^+^CD25^+^Foxp3^+^ T lymphocytes in MLN and SPL.

Analyzing B220^+^CD3^-^CD27^+^ B lymphocytes in MLN, NC (20.13±3.52) and EXP-1 (22.32±1.01) showed significantly lower titers (p=0.01) than EXP-2 (27.06±0.90), which were not statistically different (p=0.95) from one another. Similarly, in SPL, NC (38.55±3.63) and EXP-1 (41.96±2.08) showed significantly lower titers (p<0.001) than EXP-2 (55.01±1.61), which were not statistically different (p=0.63) from one another (Figure 4).

**Figure 4.**
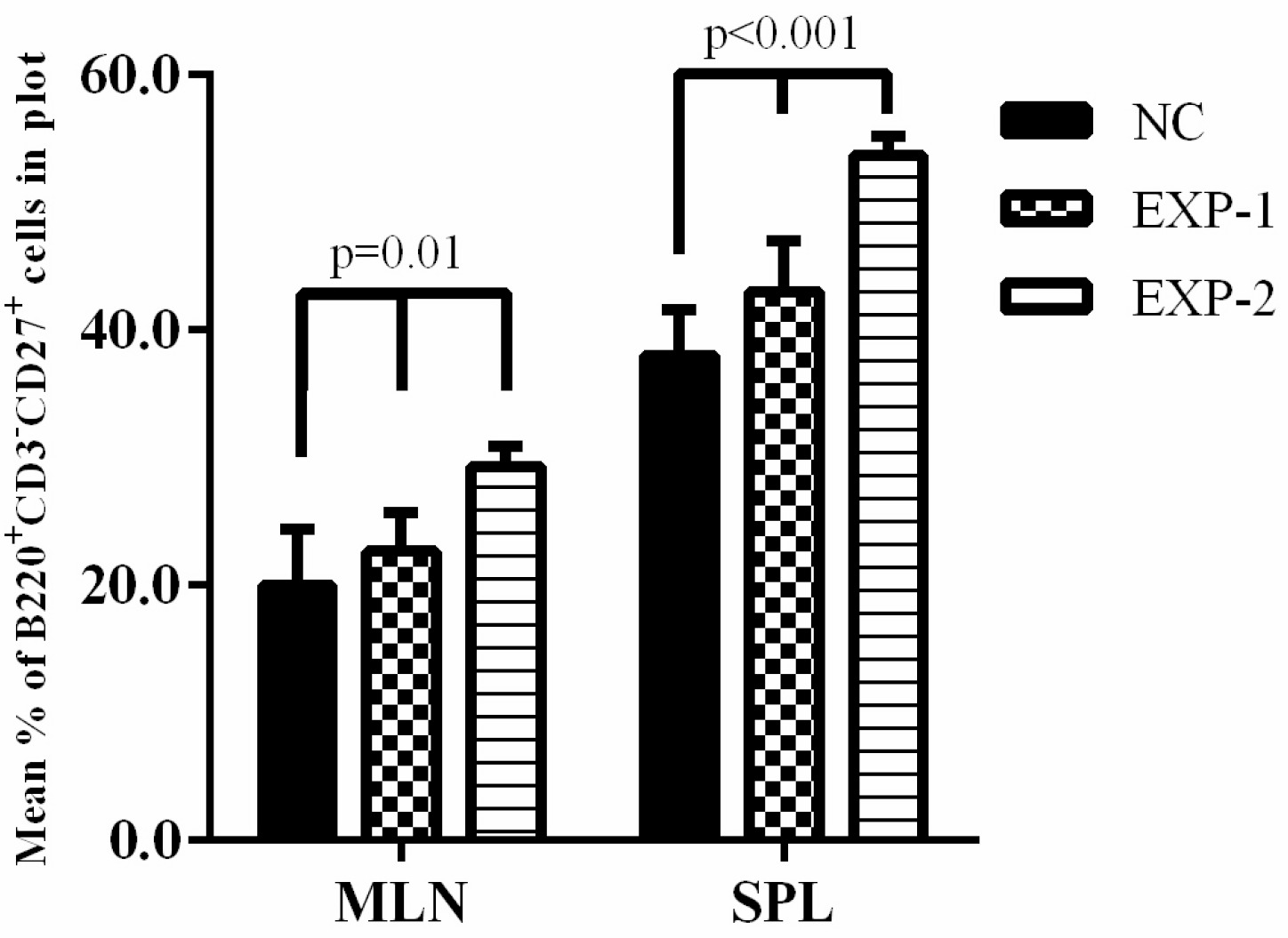
Mean group percentage in gate P3 ± SD of B220^+^CD3^-^CD27^+^ B lymphocytes in MLN and SPL.

#### 3.1.3. Histopathological Evaluation of Intestinal Samples

Gut histopathological changes were only observed in EXP-2 animals and were restricted to the duodenum. The large intestine remained intact, with no identifiable changes among the groups after preliminary macroscopic and microscopic evaluation (Images not shown). In NC and EXP-1, the mucous layer was intact and had a preserved epithelial lining along the finger-like villi. Lamina propria and epithelium had no signs suggestive of edema. EXP-1 showed rare intraepithelial mononuclear leukocytes and absence of heterotypic lymphoid clusters (Figure 5A and 5B).

**Figure 5.**
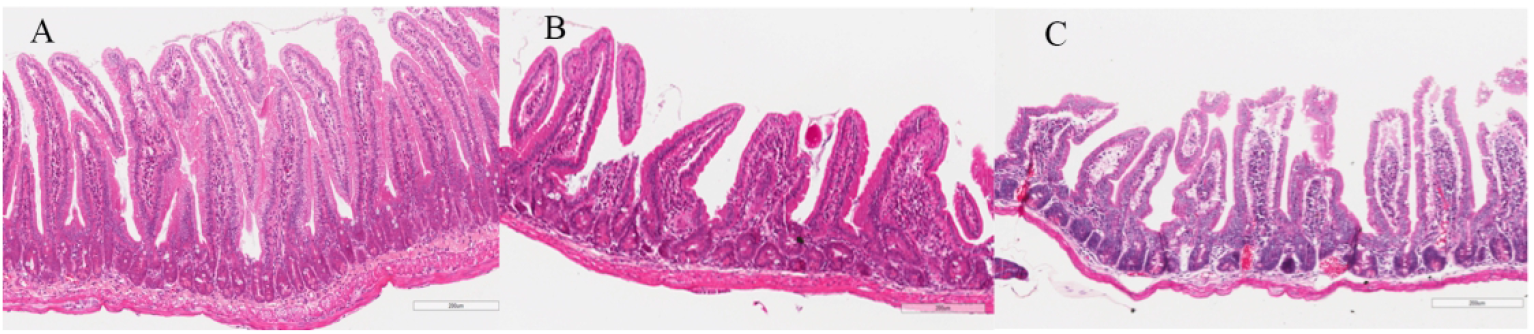
Representative images of the small intestine stained with HE shows the intestinal milieu after CD. **A**: animal from NC group showed an intestinal mucosa staged as normal. **B**: Animal from EXP-1 also showed an intestinal mucosa staged as normal. **C**: Animal from EXP-2 showed a duodenum staged as Partial Destruction. Images were captured using the ImageScope^®^ software with 20x zoom.

In EXP-2, lesions in the small intestine occurred mainly in the mucosa, without significant involvement of the submucosa, the muscular layer, or the serosa (Figure 5C). The presence of granulomas, fibrosis or fistulas was not observed. The mucous layer presented loss of integrity and continuity of its surface, due to partial tissue destruction. Villi lost their finger-like appearance. Presence of suggestive edema within the lamina propria and between the lamina propria and the epithelial lining was observed. An increase in the intraepithelial leukocytes (IEL) was also observed. After analysis, NC (Figure 5A) and EXP-1 (Figure 5B) were classified as Normal in our system and EXP-2 (Figure 5C)was staged as Partial Destruction, which is the third level of a 5-level classification system [4].

### 3.2. Results from the Second Stage Analyzed

#### 3.2.1. Anti-peanuts polyisotypic IgG and IgG1 titers

Considering all analysis points (post feeding, primary and booster inoculations, CD, and Recovery) the intergroup analysis revealed that EXP-2.2 (4.60±1.18), EXP-2.3 (4.76±0.18) and PC (4.75±1.06) had significantly higher mean polyisotypic IgG levels (p<0.001), even during the recovery period, when compared to EXP-1.2 (1.97±0.31) and NC (1.00±0.13). EXP-1.2 also presented significantly higher mean polyisotypic anti-peanut IgG levels when compared to NC (p=0.003). EXP-2.2, EXP-2.3 and PC were not statistically different from each other (p=0.99) (Figure 6A). The same pattern was also observed in anti-peanuts IgG1 titers. EXP-2.2 (2.80±1.20), EXP-2.3 (2.87±1.10) and PC (2.93±1.22) had significantly higher mean IgG1 levels (p<0.001) even during the recovery period when compared to EXP-1.2 (0.92±0.09) and NC (0.58±0.04), which were not statistically different from each other (p=0.60). Similarly, EXP-2.2, EXP-2.3 and PC were not statistically different from each other (p=0.56) (Figure 6B).

**Figure 6.**
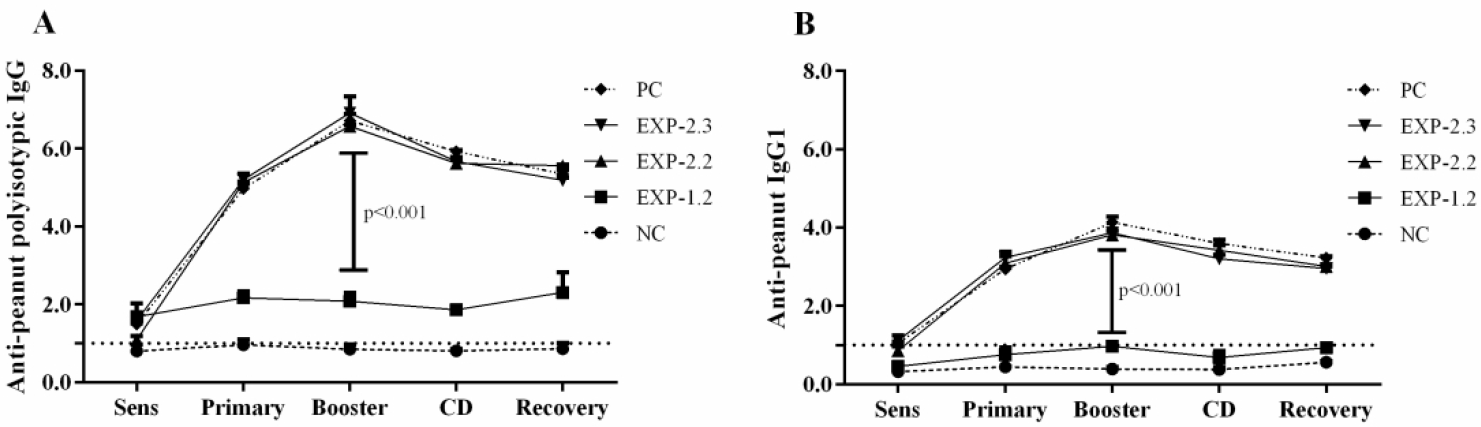
**A**) Mean group polyisotypic IgG. EXP-2.2, EXP-2.3 and PC showed significant higher titers when compared to EXP-1.2 and NC, which was immunized with saline. **B**) Mean group IgG1. EXP-2.2, EXP-2.3 and PC showed significantly higher titers when compared to EXP-1.2 and NC, which was immunized with saline. Results were expressed as ELISA index (EI), where values of EI *>*1.0 (horizontal dashed line) were considered positive. No pooled sera were used, each individual mouse was analyzed.

#### 3.2.2. Variation of effector and regulatory immune cells in MLN and SPL

After gating the lymphocyte region in MLN samples, we observed that EXP-2.3 (31.33±4.02) and PC [30.66±2.51]) showed statistically higher titers (p=0.01) compared to the other groups (NC [13.66±2.05]; EXP-1.2 [18.34±3.03]; EXP-2.2 [26.89±1.52]) regarding CD4^+^ T cells (Figure 7A). In SPL, PC (21.70±1.78) showed significantly higher titers (p=0.02) when compared to NC (10.66±1.55), EXP-1.2 (14.12±1.01), EXP-2.2 (17.22±1.10) and EXP-2.3 (19.56±1.23) (Figure 7A). Representative plots of each group can be found on supplemental figure 2.

**Figure 7.**
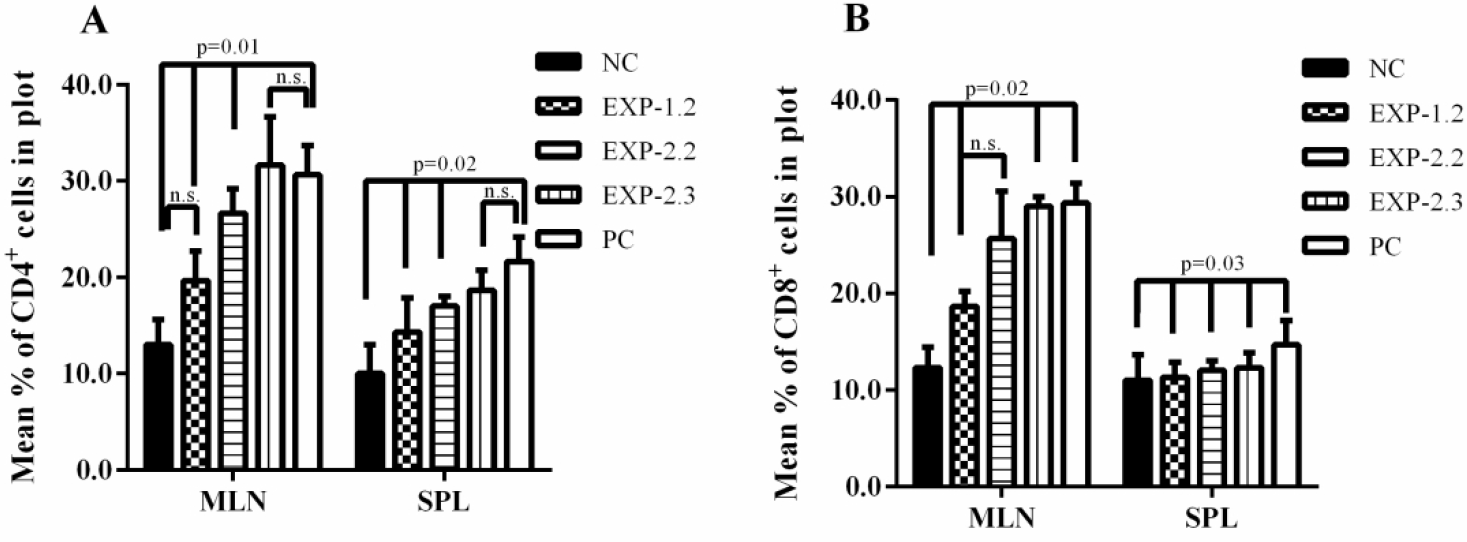
**A**) Mean plot percentage of CD4^+^ T lymphocytes in MLN and SPL in gate P3 ± SD per group. **B**) Mean plot percentage of CD8^+^ T lymphocytes in MLN and SPL in gate P3 ± SD per group.

When CD8^+^ T cells were analyzed in MLN, no statistically significant differences were observed among NC (12.33±3.88), EXP-1.2 (18.69±2.34) and EXP-2.2 (24.13±3.15) (p=0.52) (Figure 7B). EXP-2.3 (28.65±1.19) and PC (30.38±2.55) showed significant higher titers compared to NC and EXP-1.2 (p=0.02). In SPL, PC (14.89±1.06) showed significantly higher titers (p=0.03) than the other groups: NC (11.12±0.54), EXP-1.2 (11.33±0.87), EXP-2.2 (12.21±0.47), EXP-2.3 (13.34±0.91) (Figure 7B).

Regarding CD4^+^CD25^+^Foxp3^+^ T regulatory cells in MLN, NC (5.66±1.54) showed statistically lower titers (p=0.02) compared to EXP-1.2 (15.87±2.21), EXP-2.2 (13.98±1.33) and EXP-2.3 (12.33±0.99). PC (12.09±0.87) also showed statistically higher titers (p=0.01) than NC. (Figure 8A). In SPL, NC (4.32±1.56) showed significantly lower titers (p=0.001) than all the other groups [EXP-1.2 (14.06±1.08), EXP-2.2 (12.32±2.28), EXP-2.3 (11.61±2.02) and PC (13.12±2.11)] (Figure 8A).

**Figure 8.**
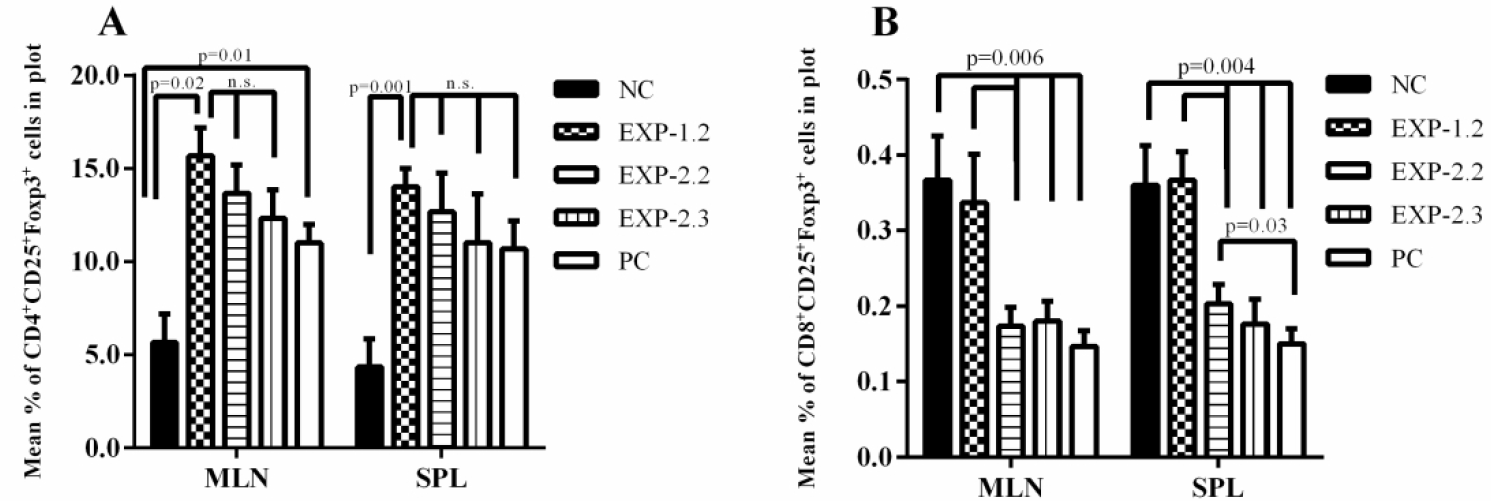
**A**) Mean plot percentage of CD4^+^CD25^+^Foxp3^+^ T lymphocytes in MLN and SPL in gate P3 ± SD per group. **B**) Mean plot percentage of CD8^+^CD25^+^Foxp3^+^ T lymphocytes in MLN and SPL in gate P3 ± SD per group.

Considering CD8^+^CD25^+^Foxp3^+^ T regulatory cells in MLN, NC (0.37±0.05) and EXP-1.2 (0.33±0.05) presented significantly higher titers (p=0.006) than EXP-2.2 (0.17±0.04), EXP-2.3 (0.18±0.04) and PC (0.14±0.02) (Figure 8B). In SPL, NC (0.36±0.05) and EXP-1.2 (0.37±0.02) showed significantly higher titers (p=0.004) than EXP-2.2 (0.20±0.02), EXP-2.3 (0.17±0.02) and PC (0.15±0.02). EXP-2.2 also showed significantly higher titers than PC (p=0.03) (Figure 8B).

When we evaluated B220^+^CD3^-^CD27^+^ memory B lymphocytes in MLN, no significant differences (p=0.17) were observed among the groups (NC [18.61±2.91]; EXP-1.2 [23.43±3.02]; EXP-2.2 [22.54±2.32]; EXP-2.3 [22.33±3.05]; and PC [18.87±3.17]) (Figure 9). In SPL, PC (54.23±3.56) showed significantly higher titers (p=0.01) than NC (36.21±3.07), EXP-1.2 (36.18±5.88), EXP-2.2 (42.12±3.11) and EXP-2.3 (43.56±1.78), which were not statistically different (p=0.11) from one another. (Figure 9).

**Figure 9.**
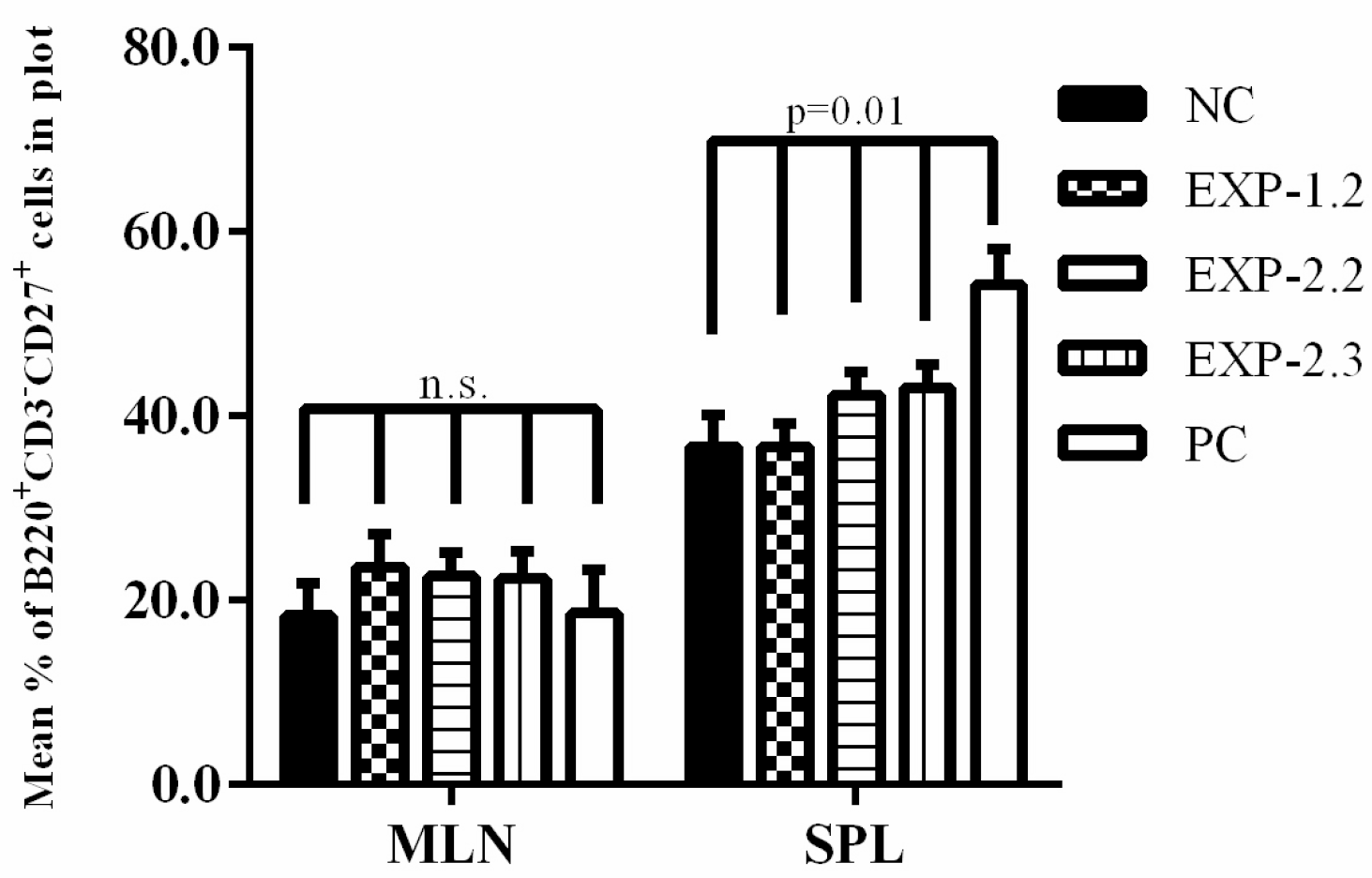
Mean plot percentage of B220^+^CD3^-^CD27^+^ memory B cells in MLN and SPL in gate P3 ± SD per group.

#### 3.2.3. Histopathological Evaluation of Intestinal Samples

NC showed preservation of all intestinal layers, as well as the same pattern of cellularity in lamina propria and lining epithelium. It was staged as Normal in the histomorphometric classification system developed by our group (Figure 10A).

**Figure 10.**
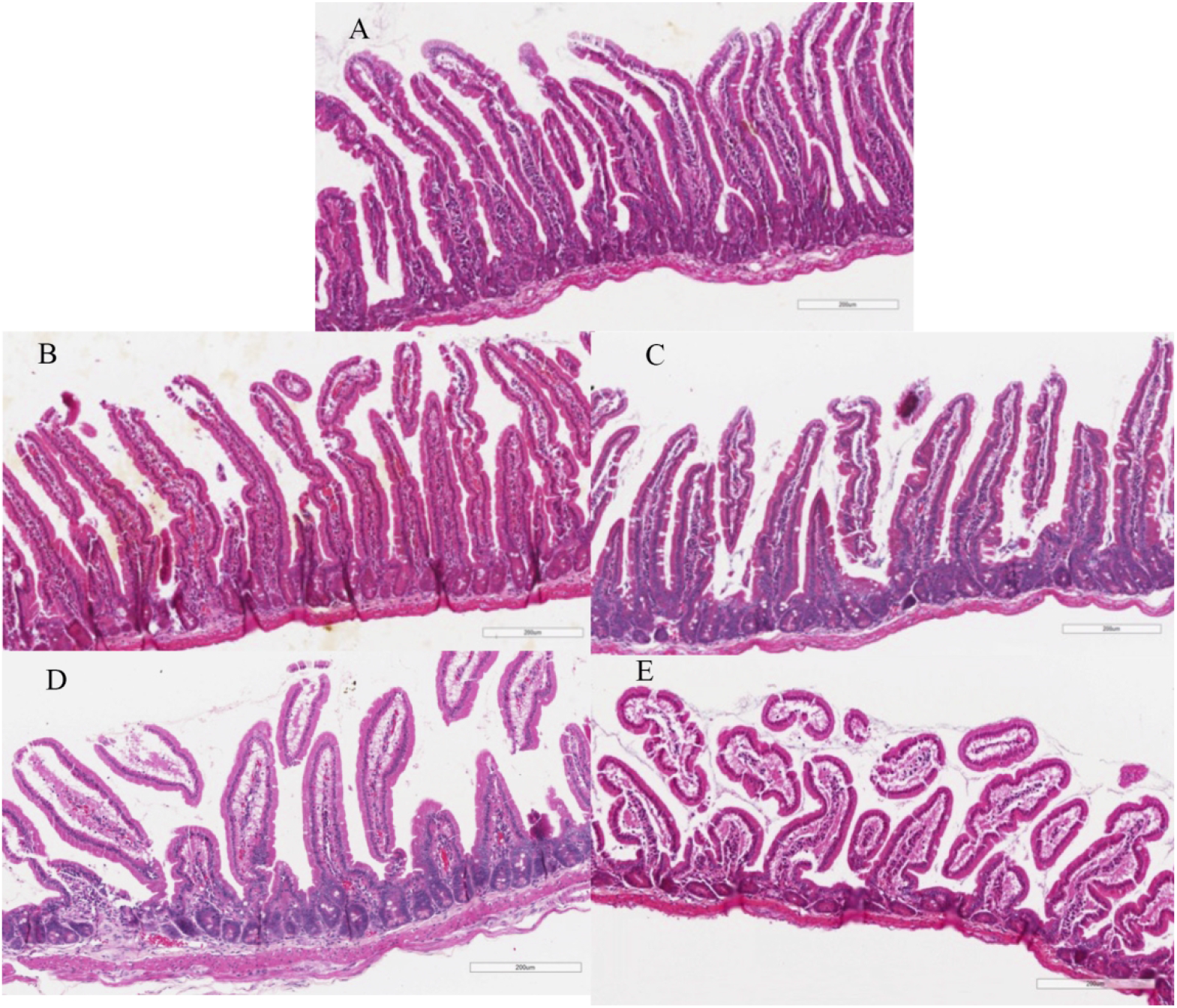
Representative images of the small intestine stained with HE show the intestinal milieu after CD. **A**: animal from NC group showed an intestinal mucosa staged as normal. **B**: Animal from EXP-1.2 also showed an intestinal mucosa staged as normal. **C**: Animal from EXP-2.2 showed a duodenum staged as Infiltrative. **D and E**: Animals from EXP-2.3 and PC showed duodenum staged as Partial Destruction. Images were captured using the ImageScope^®^ software with 20x zoom.

EXP-1.2 presented a mucosal layer with an intact and preserved epithelium and finger-like villi. Lamina propria and epithelium without signs suggestive of edema, presence of an increase in intraepithelial mononuclear leukocytes, greater than in normal animals and significantly less than in inflamed animals. We also did not find the presence of heterotypic lymphoid clusters. This group was also staged as Normal (Figure 10B).

EXP-2.2 showed similarity with EXP-1.2, both from a morphological and cellular point of view. This group, which received hybrid diet, did not develop the inflammatory process, and was protected when we evaluated the morphological characteristics, with increased intraepithelial infiltrate. EXP-2.2 animals were classified in the Infiltrative stage (Figure 10C). In EXP-2.3 and PC, the changes observed were restricted to the duodenum. The large intestine remained intact, without identifiable changes similar to NC, after preliminary macroscopic and microscopic evaluation. Lesions in the small intestine occurred mainly in the mucosa, without significant involvement of the submucosa, the muscular layer or the serosa, and the presence of granulomas, fibrosis or fistulas was not observed. The evaluation of the mucosal layer showed loss of integrity and continuity of its surface, due to tissue destruction, which in most cases is partial and not total. The villi lost their finger-like appearance. Presence of suggestive signs of edema within the lamina propria and between the lamina propria and the lining epithelium also occurred. Both groups were staged as Partial Destruction (Figure 10D and 10E).

## 4. Discussion

In this study we pondered whether antigen-specific memory-cell activation can boost oral tolerance and modulate the recovery kinetics of the inflamed intestinal mucosa. A scrutiny in the literature showed that the clinical symptoms during food allergic events vary according to eating habits and food components [36, 37]. Clinical conditions of the animals through body weight, food consumption and signs of anaphylactic reactions were assessed during all stages of the experiment. All groups showed weight gain consistent with growth and with the handling procedures to which they were submitted and with the characteristics of the C57Bl/6 lineage, as observed in previous works from our group [33, 38] and in the literature [39, 40]. Even during the offering of a challenge food containing exclusively peanuts, no signs of anaphylactic reactions were visible nor were there significant alterations of the feces consistency in any group. The introduction of a CD composed exclusively of the allergen prevents animals from avoiding the allergenic food, opposed to what happens when animals have free choice between allergenic and other food sources [29].

Studies have shown that food allergies may develop even with low titers or absence of IgE antibodies [41-43], thus leading to the current agreement in the literature that not all food allergies are exclusively caused by IgE [44-46]. Here, Morita et al. classification method was used: 1) “IgE-mediated,” 2) “non-IgE-mediated and 3) “combined IgE- and cell-mediated” [47]. Here, using C57BL/6 mice, no peanut-specific IgE was found (data not shown), thus correlating to a Non–IgE-mediated food allergy [48, 49]. Furthermore, other studies from our group have shown that our mouse model does not induce IgE in C57BL/6 mice, only in Balb/c mice [50]. Studies in the literature have also revealed the lack of IgE-mediated anaphylactic reactions in peanut-sensitized C57BL/6 mice [51, 52].

Regarding anti-peanuts polyisotypic IgG, NC, EXP-1, and EXP-1.2 had significantly lower titers compared to all other groups. This was already expected, as these groups either did not receive peanuts (NC) or were submitted to the tolerance induction protocol to peanuts (EXP-1 and EXP-1.2) through feeding, prior to immunization [53]. Conversely, high IgG titers were observed in the groups submitted to systemic immunization followed by a CD to induce intestinal inflammation. This result is similar to the clinical scenario observed in allergic children. A frequent food allergy characteristic is an inflammatory reaction in the gut when allergic patients ingest the offending food [54]. However, the presence of high antibody titers is not enough to predict an adverse reaction to food, as stated by the Canadian Society of Allergy and Clinical Immunology, since not all individuals with high antibody levels to food will present adverse reactions upon consumption [55].

Serum specific IgG1 is considered an indicator of a Th2 type immune response, hence a marker for food allergy [56, 57], for their ability to drive anaphylaxis through binding to FcγRIII expressed on macrophages or basophils’ surfaces, that ultimately leads to Platelet-Activating Factor (PAF) release [58, 59]. Consequently, the IgG subclass IgG1 was also assessed in our work. In a similar manner as observed in polyisotypic IgG titers, NC, EXP-1 and EXP-1.2 showed lower serum anti-peanut IgG1 titers than the EXP-2 group and subgroups. As stated before, no clinical alterations were observed during CD.

Our results agree to prior findings of other well studied proteins such as OVA (64,77). The offer of peanuts in the mouse diet, can either induce tolerance or sensitization depending on the intestinal context. A common finding in tolerance induction is significantly lower polyisotypic IgG titers to the tolerogen, however higher than those who never encountered the allergen before. On the other hand, in tolerant animals, anti-peanut IgG1 is consistently as low as those in normal controls.

To better understand the characteristics of the immune response, both local and systemic, phenotypic analysis of cells in MLN and SPL were performed after CD and after the recovery period. We evaluated populations of effector (CD4^+^ and CD8^+^), regulatory T cells (CD4^+^CD25^+^Foxp3^+^ and CD8^+^CD25^+^Foxp3^+^), and of memory B cells (B220^+^CD3^-^CD27^+^). These cell types were chosen because they are hallmarks of effector and regulatory T cells populations respectively. These are the two major populations present in allergic and tolerance reactions [60, 61], along with memory B cells [62, 63].

Considering that all experiments were terminated during CD (50% on peanut CD and 50% on OVA CD), we expected that the profile of immunological cells in EXP-2 would be different from NC and EXP-1 in the local draining lymph nodes. A significant increase in both CD4^+^ and CD8^+^ T cells was observed in EXP-2 in MLN, however, the systemic cell populations as measured in the spleen showed an increase in CD8^+^ T cells, but not in CD4^+^ T cells. The increase of both CD4^+^ and CD8^+^ T cells in MLN is probably due to the activation of these cells by dietary antigens, in the gut of the immunized (allergic) animals. This result correlates to the increased IEL observed in the duodenum and is in accordance with the literature [64]. The increment of T cell population is not exclusively due to the introduction of peanuts in the diet. Animals receiving the diet for the first time or as a challenge food after being submitted to the tolerance protocol (which includes systemic immunization) do not alter both CD4^+^ and CD8^+^ T cell populations in the draining lymph nodes. IFN-γ secreted by CD8^+^ T lymphocytes acts as a suppressor of antigen specific CD4^+^ Th2 cells in mouse models using OVA. CD8^+^ T lymphocytes also secrete TGF-ß and IL-10 both suppressor molecules of inflammatory reaction. The balance of the cytokine network in the mucosal *milleu* is not an antigen-specific immune response although initially activated by antigens. [65, 66].

The increase in CD8^+^ T cells in EXP-2 and PC in SPL can be explained by feedback responses in chronic inflammatory processes. Analyzing the percentage of CD4^+^CD25^+^Foxp3^high^ regulatory T lymphocytes, also called thymic regulatory T cells [67], in both MLN and SPL, showed an increase in EXP-1 and EXP-1.2, which correlates to the induction of oral tolerance in the literature [68]. These cells also play an important role in preserving tissue homeostasis [69]. We also observed an increase of CD8^+^CD25^+^Foxp3^+^ regulatory T lymphocytes in EXP-1 and EXP-1.2 and a decrease in EXP-2.2, EXP-2.3, and PC in both MLN and SPL. This population is poorly studied and is usually a heterogeneous group [70, 71]. Since CD8^+^ T cells bind to MHC-I, CD8^+^ regulatory T lymphocytes are more studied in transplantation mouse models in comparison to food allergy, however some studies report that CD8^+^ CD25^+^Foxp3^+^ T cells can ameliorate inflammatory responses in allergic diseases [71, 72]. These TGF-ß expressing regulatory T cells are crucial for the induction and maintenance of oral tolerance together with CD4^+^ regulatory T cells [73]. This suppressor activity can explain the results stated above.

B220^+^CD3^-^CD27^+^ memory B cells were also assessed in our study. These antigen-specific memory cells were first described over 20 years ago and are responsible for quick response upon antigen restimulation by switching to a B220^low^ phenotype [74, 75]. This may indicate that EXP-2.3 was sensitized to OVA during the CD, which has been observed in previous studies in our group [32] and also in the literature [76, 77]. B220^+^CD3^-^CD27^+^ memory B cells are also noteworthy for induction of class-switching to IgG1 in plasma cells through activation of Th2 memory T cells [78], which can explain why EXP-2 and PC showed an increase of this population. Together with the normal duodenum parameters, the B220^+^CD3^-^ CD27^+^ memory B cells titers in EXP-2.2 can indicate that this group was protected against inflammation in the gut by the activation of the memory cells generated when animals were offered mouse chow for the first time in the oral sensitization phase. As these animals went from an inflammatory milieu in the gut during the CD to a more tolerogenic milieu in the recovery period, it is possible that a bystander effect occurred, where the memory B cells and the regulatory T cells induced a change of the Th2-type response to a more tolerogenic type response, as previously described in adoptive transfer studies [79, 80].

It was imperative to examine the gut of the animals due to the CD phase. Our analysis showed that the gut histological profile correlate to the findings from the serum, MLN, and SPL of the animals. Those with higher IgG and IgG1 titers (sensitization to peanuts) presented an inflamed gut along with higher effector T cells [81]. A classification system for mouse inflamed duodenum developed by our group was used to easily compare different protocols [35]. EXP-2.3 showed a partially destructed duodenum, which may indicate that this group was also sensitized to OVA, although specific anti-OVA antibodies were not the focus of this work. The partial destruction stage in which EXP-2, EXP-2.3, and PC were classified, describe progressive alterations of the mucosal architecture with an increasing number of the leukocytes within the epithelial layer, therefore presenting hypertrophy of the lamina propria, as well as depletion of goblet cells. Reduction of these cells has an important impact on mucosal integrity due to the proportional decrease in mucin secretion, with consequent impairment of the protective function of the mucus [82]. This goblet cell reduction was also observed in the previously developed gut inflammation classification systems [83-85] and in ulcerative colitis studies in mice [86].

Although there was a time lapse between CD and the end of the experiment (recovery period), the perpetuation of an inflammatory process in the gut correlates with the decrease in regulatory T cells in EXP-2 and PC. This can be justified by an accumulation of extracellular ATP (adenosine 5’triphosphate) when tissue damage occurs. These molecules bind to the P2X7R ligand and promotes cell death in some regulatory T cells, while others convert into a Th17 phenotype [87].

Other studies still need to be performed to comprehend the role of the various immune cell types as well as cytokines and immunoglobulins involved in this process through time. For instance, IgA evaluation was not possible during this work, but previous data from our group as well as in the literature suggest this immunoglobulin may play a significant role in our mouse model [88]. Here, we confirmed that a bystander suppression may occur when the animals were offered the allergenic food associated to the ingestion of tolerogenic food (mouse chow). In conclusion, when a tolerogenic food is added to a diet containing allergen, it can ameliorate the induction of local inflammation, as well as increase mucosa recovery by a bystander effect, possibly because of regulatory cells expansion and previously induced memory cells activation.

## Acknowledgements

The authors would like to thank Professor Maira Platais for the English review.

## References

[1] Lee T, Edwards-Salmon S, Vickery BP. Current and future treatments for peanut allergy. Clin Exp Allergy 2023;53:10–24. doi: 10.1111/cea.14244

[2] Strid J, Thomson M, Hourihane J, Kimber I, Strobel S. A novel model of sensitization and oral tolerance to peanut protein. Immunology 2004;113:293–303. doi: 10.1111/j.1365-2567.2004.01989.x

[3] Reshma A, Baranwal AK. Child with Allergies or Allergic Reactions. Indian journal of pediatrics 2017;85:60–5. doi: 10.1007/s12098-017-2436-8

[4] Pereira e Silva A, Soares JRA, Mattos EBA, Josetti C, Mazza-Guimarães I, Campos SM, et al. A histomorphometric classification system for normal and inflamed mouse duodenum— Quali - quantitative approach. Int J Exp Pathol 2018;00:1–10. doi: https://doi.org/10.1111/iep.12286

[5] Arasi S, Castagnoli R, Pajno GB. Oral immunotherapy in pediatrics. Pediatric allergy and immunology : official publication of the European Society of Pediatric Allergy and Immunology 2020;31 Suppl 24:51–3. doi: 10.1111/pai.13159

[6] Bartnikas LM, Huffaker MF, Sheehan WJ, Kanchongkittiphon W, Petty CR, Leibowitz R, et al. Racial and socioeconomic differences in school peanut-free policies. The journal of allergy and clinical immunology In practice 2020;8:340–2 e1. doi: 10.1016/j.jaip.2019.06.036

[7] Skypala IJ. Food-Induced Anaphylaxis: Role of Hidden Allergens and Cofactors. Frontiers in immunology 2019;10:673. doi: 10.3389/fimmu.2019.00673

[8] Cherkaoui S, Ben-Shoshan M, Alizadehfar R, Asai Y, Chan E, Cheuk S, et al. Accidental exposures to peanut in a large cohort of Canadian children with peanut allergy. Clinical and translational allergy 2015;5:16. doi: 10.1186/s13601-015-0055-x

[9] Muraro A, Sublett JW, Haselkorn T, Nilsson C, Casale TB. Incidence of anaphylaxis and accidental peanut exposure: A systematic review. Clinical and translational allergy 2021;11:e12064. doi: 10.1002/clt2.12064

[10] Jones SM, Kim EH, Nadeau KC, Nowak-Wegrzyn A, Wood RA, Sampson HA, et al. Efficacy and safety of oral immunotherapy in children aged 1-3 years with peanut allergy (the Immune Tolerance Network IMPACT trial): a randomised placebo-controlled study. Lancet 2022;399:359–71. doi: 10.1016/S0140-6736(21)02390-4

[11] Grzeskowiak LE, Tao B, Aliakbari K, Chegeni N, Morris S, Chataway T. Oral immunotherapy using boiled peanuts for treating peanut allergy: An open-label, single-arm trial. Clin Exp Allergy 2023. doi: 10.1111/cea.14254

[12] Gupta RS, Lau CH, Sita EE, Smith B, Greenhawt MJ. Factors associated with reported food allergy tolerance among US children. Annals of allergy, asthma & immunology : official publication of the American College of Allergy, Asthma, & Immunology 2013;111:194–8 e4. doi: 10.1016/j.anai.2013.06.026

[13] Nitsche C, Westerlaken-van Ginkel CD, Kollen BJ, Sprikkelman AB, Koppelman GH, Dubois AEJ. Eliciting dose is associated with tolerance development in peanut and cow’s milk allergic children. Clinical and translational allergy 2019;9:58. doi: 10.1186/s13601-019-0298-z

[14] Hourihane JO, Roberts SA, Warner JO. Resolution of peanut allergy: case-control study. BMJ 1998;316:1271–5. doi: 10.1136/bmj.316.7140.1271

[15] Hourihane JO, Allen KJ, Shreffler WG, Dunngalvin G, Nordlee JA, Zurzolo GA, et al. Peanut Allergen Threshold Study (PATS): Novel single-dose oral food challenge study to validate eliciting doses in children with peanut allergy. The Journal of allergy and clinical immunology 2017;139:1583–90. doi: 10.1016/j.jaci.2017.01.030

[16] Monian B, Tu AA, Ruiter B, Morgan DM, Petrossian PM, Smith NP, et al. Peanut oral immunotherapy differentially suppresses clonally distinct subsets of T helper cells. The Journal of clinical investigation 2022;132. doi: 10.1172/JCI150634

[17] Vazquez-Cortes S, Jaqueti P, Arasi S, Machinena A, Alvaro-Lozano M, Fernandez-Rivas M. Safety of Food Oral Immunotherapy: What We Know, and What We Need to Learn. Immunology and allergy clinics of North America 2020;40:111–33. doi: 10.1016/j.iac.2019.09.013

[18] Zhang W, Dhondalay GK, Liu TA, Kaushik A, Hoh R, Kwok S, et al. Gastrointestinal gammadelta T cells reveal differentially expressed transcripts and enriched pathways during peanut oral immunotherapy. Allergy 2022. doi: 10.1111/all.15250

[19] Nowak-Wegrzyn A, Berin MC, Mehr S. Food Protein-Induced Enterocolitis Syndrome. The journal of allergy and clinical immunology In practice 2020;8:24–35. doi: 10.1016/j.jaip.2019.08.020

[20] Cao S, Nagler CR. Interpreting success or failure of peanut oral immunotherapy. The Journal of clinical investigation 2022;132. doi: 10.1172/JCI155255

[21] Investigators PGoC, Vickery BP, Vereda A, Casale TB, Beyer K, du Toit G, et al. AR101 Oral Immunotherapy for Peanut Allergy. The New England journal of medicine 2018;379:1991–2001. doi: 10.1056/NEJMoa1812856

[22] Chinthrajah RS, Purington N, Andorf S, Long A, O’Laughlin KL, Lyu SC, et al. Sustained outcomes in oral immunotherapy for peanut allergy (POISED study): a large, randomised, double-blind, placebo-controlled, phase 2 study. Lancet 2019;394:1437–49. doi: 10.1016/S0140-6736(19)31793-3

[23] Kilkenny C, Browne WJ, Cuthill IC, Emerson M, Altman DG. Improving bioscience research reporting: the ARRIVE guidelines for reporting animal research. PLoS biology 2010;8:1–5. doi: 10.1371/journal.pbio.1000412

[24] Whipple B, Agar J, Zhao J, Pearce DA, Kovacs AD. The acidified drinking water-induced changes in the behavior and gut microbiota of wild-type mice depend on the acidification mode. Scientific reports 2021;11:2877. doi: 10.1038/s41598-021-82570-0

[25] Tanner RS, James SA. Rapid bactericidal effect of low pH against Pseudomonas aeruginosa. J Ind Microbiol 1992;10:229–32

[26] Larsen JM, Bogh KL. Animal models of allergen-specific immunotherapy in food allergy: Overview and opportunities. Clin Exp Allergy 2018;48:1255–74. doi: 10.1111/cea.13212

[27] Kreuter R, Wankell M, Ahlenstiel G, Hebbard L. The role of obesity in inflammatory bowel disease. Biochim Biophys Acta Mol Basis Dis 2019;1865:63–72. doi: 10.1016/j.bbadis.2018.10.020

[28] Reeves PG, Nielsen FH, Fahey GC, Jr. AIN-93 purified diets for laboratory rodents: final report of the American Institute of Nutrition ad hoc writing committee on the reformulation of the AIN-76A rodent diet. The Journal of nutrition 1993;123:1939–51. doi: 10.1093/jn/123.11.1939

[29] Teixeira G, Paschoal PO, de Oliveira VL, Pedruzzi MM, Campos SM, Andrade L, et al. Diet selection in immunologically manipulated mice. Immunobiology 2008;213:1–12

[30] Mirotti L, Mucida D, de Sa-Rocha Lc, Costa-Pinto FA, Russo M. Food aversion: a critical balance between allergen-specific IgE levels and taste preference. Brain, behavior, and immunity 2010;24:370–5. doi: 10.1016/j.bbi.2009.12.006

[31] Costa-Pinto FA, Basso AS. Neural and behavioral correlates of food allergy. Chem Immunol Allergy 2012;98:222–39. doi: 10.1159/000336525

[32] Pereira ESA, Marmello BO, Soares JRA, Mazza-Guimaraes I, Teixeira G. Induction of food tolerance is dependent on intestinal inflammatory state. Immunol Lett 2021;234:33–43. doi: 10.1016/j.imlet.2021.04.009

[33] Campos SM, de Oliveira VL, Lessa L, Vita M, Conceicao M, Andrade LA, et al. Maternal immunomodulation of the offspring’s immunological system. Immunobiology 2014;219:813–21. doi: 10.1016/j.imbio.2014.07.001

[34] Sadtler K, Elisseeff JH. Analyzing the scaffold immune microenvironment using flow cytometry: practices, methods and considerations for immune analysis of biomaterials. Biomater Sci 2019;7:4472–81. doi: 10.1039/c9bm00349e

[35] Pereira ESA, Soares JRA, Mattos EBA, Josetti C, Guimaraes IM, Campos SMN, et al. A histomorphometric classification system for normal and inflamed mouse duodenum-Quali-quantitative approach. Int J Exp Pathol 2018;99:189–98. doi: 10.1111/iep.12286

[36] Tedner SG, Asarnoj A, Thulin H, Westman M, Konradsen JR, Nilsson C. Food allergy and hypersensitivity reactions in children and adults-A review. J Intern Med 2022;291:283–302. doi: 10.1111/joim.13422

[37] Sicherer SH, Abrams EM, Nowak-Wegrzyn A, Hourihane JO. Managing Food Allergy When the Patient Is Not Highly Allergic. The journal of allergy and clinical immunology In practice 2022;10:46–55. doi: 10.1016/j.jaip.2021.05.021

[38] Paschoal PO, Campos SM, Pedruzzi MM, Garrido V, Bisso M, Antunes DM, et al. Food allergy/hypersensitivity: antigenicity or timing? Immunobiology 2009;214:269–78. doi: 10.1016/j.imbio.2008.09.007

[39] Cardoso CR, Teixeira G, Provinciatto PR, Godoi DF, Ferreira BR, Milanezi CM, et al. Modulation of mucosal immunity in a murine model of food-induced intestinal inflammation. Clin Exp Allergy 2008;38:338–49

[40] Tordesillas L, Berin MC. Mechanisms of Oral Tolerance. Clin Rev Allergy Immunol 2018;55:107–17. doi: 10.1007/s12016-018-8680-5

[41] Lee E, Barnes EH, Mehr S, Campbell DE. Differentiating Acute Food Protein-Induced Enterocolitis Syndrome From Its Mimics: A Comparison of Clinical Features and Routine Laboratory Biomarkers. The journal of allergy and clinical immunology In practice 2019;7:471–8 e3. doi: 10.1016/j.jaip.2018.10.020

[42] Shakoor Z, AlFaifi A, AlAmro B, AlTawil LN, AlOhaly RY. Prevalence of IgG-mediated food intolerance among patients with allergic symptoms. Ann Saudi Med 2016;36:386–90. doi: 10.5144/0256-4947.2016.386

[43] Wang HY, Li Y, Li JJ, Jiao CH, Zhao XJ, Li XT, et al. Serological investigation of IgG and IgE antibodies against food antigens in patients with inflammatory bowel disease. World J Clin Cases 2019;7:2189–203. doi: 10.12998/wjcc.v7.i16.2189

[44] Johansson SG, Hourihane JO, Bousquet J, Bruijnzeel-Koomen C, Dreborg S, Haahtela T, et al. A revised nomenclature for allergy. An EAACI position statement from the EAACI nomenclature task force. Allergy 2001;56:813–24. doi: all001 [pii]

[45] Muraro A, Werfel T, Hoffmann-Sommergruber K, Roberts G, Beyer K, Bindslev-Jensen C, et al. EAACI food allergy and anaphylaxis guidelines: diagnosis and management of food allergy. Allergy 2014;69:1008–25. doi: 10.1111/all.12429

[46] Fiocchi A, Bognanni A, Brozek J, Ebisawa M, Schunemann H, group WDg. World Allergy Organization (WAO) Diagnosis and Rationale for Action against Cow’s Milk Allergy (DRACMA) Guidelines update - I - Plan and definitions. The World Allergy Organization journal 2022;15:100609. doi: 10.1016/j.waojou.2021.100609

[47] Morita H, Nomura I, Matsuda A, Saito H, Matsumoto K. Gastrointestinal food allergy in infants. Allergology international : official journal of the Japanese Society of Allergology 2013;62:297–307. doi: 10.2332/allergolint.13-RA-0542

[48] Nowak-Wegrzyn A, Katz Y, Mehr SS, Koletzko S. Non-IgE-mediated gastrointestinal food allergy. The Journal of allergy and clinical immunology 2015;135:1114–24. doi: 10.1016/j.jaci.2015.03.025

[49] Okazaki F, Wakiguchi H, Korenaga Y, Takahashi K, Yasudo H, Fukuda K, et al. Food Protein-Induced Enterocolitis Syndrome in Children with Down Syndrome: A Pilot Case-Control Study. Nutrients 2022;14. doi: 10.3390/nu14020388

[50] Soares JRA, Pereira ESA, de Souza Oliveira AL, Guimaraes IM, das Neves Faccini CRJ, de Aquino Mattos EB, et al. Allergen extraction: Factors influencing immunogenicity and sensitivity of immunoassays. J Immunol Methods 2021;498:113125. doi: 10.1016/j.jim.2021.113125

[51] Paolucci M, Homere V, Waeckerle-Men Y, Wuillemin N, Bieli D, Pengo N, et al. Strain matters in mouse models of peanut-allergic anaphylaxis: Systemic IgE-dependent and Ara h 2-dominant sensitization in C3H mice. Clin Exp Allergy 2023. doi: 10.1111/cea.14279

[52] Orgel K, Smeekens JM, Ye P, Fotsch L, Guo R, Miller DR, et al. Genetic diversity between mouse strains allows identification of the CC027/GeniUnc strain as an orally reactive model of peanut allergy. The Journal of allergy and clinical immunology 2019;143:1027–37 e7. doi: 10.1016/j.jaci.2018.10.009

[53] Teixeira GAPB, Antunes DMF, Castro Júnior ABd, Costa JPd, Paschoal PO, Campos SMN, et al. Induction of an antigen specific gut inflammatory reaction in mice and rats: a model for human inflammatory bowel disease. Brazilian Archives of Biology and Technology 2009;52:601–9

[54] Calvani M, Anania C, Cuomo B, D’Auria E, Decimo F, Indirli GC, et al. Non-IgE- or Mixed IgE/Non-IgE-Mediated Gastrointestinal Food Allergies in the First Years of Life: Old and New Tools for Diagnosis. Nutrients 2021;13. doi: 10.3390/nu13010226

[55] Carr S, Chan E, Lavine E, Moote W. CSACI Position statement on the testing of food-specific IgG. Allergy Asthma Clin Immunol 2012;8:12. doi: 10.1186/1710-1492-8-12

[56] Weidmann E, Samadi N, Klems M, Heiden D, Seppova K, Ret D, et al. Mouse Chow Composition Influences Immune Responses and Food Allergy Development in a Mouse Model. Nutrients 2018;10. doi: 10.3390/nu10111775

[57] Chen C, Lianhua L, Nana S, Yongning L, Xudong J. Development of a BALB/c mouse model for food allergy: comparison of allergy-related responses to peanut agglutinin, beta-lactoglobulin and potato acid phosphatase. Toxicol Res (Camb) 2017;6:251–61. doi: 10.1039/c6tx00371k

[58] Miyajima I, Dombrowicz D, Martin TR, Ravetch JV, Kinet JP, Galli SJ. Systemic anaphylaxis in the mouse can be mediated largely through IgG1 and Fc gammaRIII. Assessment of the cardiopulmonary changes, mast cell degranulation, and death associated with active or IgE- or IgG1-dependent passive anaphylaxis. The Journal of clinical investigation 1997;99:901–14. doi: 10.1172/JCI119255

[59] Dekkers G, Bentlage AEH, Stegmann TC, Howie HL, Lissenberg-Thunnissen S, Zimring J, et al. Affinity of human IgG subclasses to mouse Fc gamma receptors. MAbs 2017;9:767–73. doi: 10.1080/19420862.2017.1323159

[60] Stringari LL, Covre LP, da Silva FDC, de Oliveira VL, Campana MC, Hadad DJ, et al. Increase of CD4+CD25highFoxP3+ cells impairs in vitro human microbicidal activity against Mycobacterium tuberculosis during latent and acute pulmonary tuberculosis. PLoS Negl Trop Dis 2021;15:e0009605. doi: 10.1371/journal.pntd.0009605

[61] Brummelman J, Pilipow K, Lugli E. The Single-Cell Phenotypic Identity of Human CD8(+) and CD4(+) T Cells. Int Rev Cell Mol Biol 2018;341:63–124. doi: 10.1016/bs.ircmb.2018.05.007

[62] Machura E, Mazur B, Pieniazek W, Karczewska K. Expression of naive/memory (CD45RA/CD45RO) markers by peripheral blood CD4+ and CD8 + T cells in children with asthma. Archivum immunologiae et therapiae experimentalis 2008;56:55–62. doi: 10.1007/s00005-008-0005-6

[63] Jiang Q, Yang G, Liu Q, Wang S, Cui D. Function and Role of Regulatory T Cells in Rheumatoid Arthritis. Frontiers in immunology 2021;12:626193. doi: 10.3389/fimmu.2021.626193

[64] Yu W, Zhou X, Dunham D, Lyu SC, Manohar M, Zhang W, et al. Allergen-specific CD8(+) T cells in peanut-allergic individuals. The Journal of allergy and clinical immunology 2019;143:1948–52. doi: 10.1016/j.jaci.2019.01.011

[65] Dahlman-Hoglund A, Dahlgren U, Ahlstedt S, Hanson LA, Telemo E. Bystander suppression of the immune response to human serum albumin in rats fed ovalbumin. Immunology 1995;86:128–33

[66] Backstrom NF, Dahlgren UI. Bystander suppression of collagen-induced arthritis in mice fed ovalbumin. Arthritis Res Ther 2004;6:R151–60. doi: 10.1186/ar1150

[67] Giganti G, Atif M, Mohseni Y, Mastronicola D, Grageda N, Povoleri GA, et al. Treg cell therapy: How cell heterogeneity can make the difference. Eur J Immunol 2021;51:39–55. doi: 10.1002/eji.201948131

[68] Cosovanu C, Neumann C. The Many Functions of Foxp3(+) Regulatory T Cells in the Intestine. Frontiers in immunology 2020;11:600973. doi: 10.3389/fimmu.2020.600973

[69] Laukova M, Glatman Zaretsky A. Regulatory T cells as a therapeutic approach for inflammatory bowel disease. Eur J Immunol 2022:e2250007. doi: 10.1002/eji.202250007

[70] Niederlova V, Tsyklauri O, Chadimova T, Stepanek O. CD8(+) Tregs revisited: A heterogeneous population with different phenotypes and properties. Eur J Immunol 2021;51:512–30. doi: 10.1002/eji.202048614

[71] Flippe L, Bezie S, Anegon I, Guillonneau C. Future prospects for CD8(+) regulatory T cells in immune tolerance. Immunological reviews 2019;292:209–24. doi: 10.1111/imr.12812

[72] Lin L, Dai F, Wei J, Chen Z. CD8(+) Tregs ameliorate inflammatory reactions in a murine model of allergic rhinitis. Allergy Asthma Clin Immunol 2021;17:74. doi: 10.1186/s13223-021-00577-8

[73] Mishra S, Liao W, Liu Y, Yang M, Ma C, Wu H, et al. TGF-beta and Eomes control the homeostasis of CD8+ regulatory T cells. The Journal of experimental medicine 2021;218. doi: 10.1084/jem.20200030

[74] McHeyzer-Williams LJ, Cool M, McHeyzer-Williams MG. Antigen-specific B cell memory: expression and replenishment of a novel b220(-) memory b cell compartment. The Journal of experimental medicine 2000;191:1149–66. doi: 10.1084/jem.191.7.1149

[75] Kodali S, Li M, Budai MM, Chen M, Wang J. Protection of Quiescence and Longevity of IgG Memory B Cells by Mitochondrial Autophagy. Journal of immunology 2022;208:1085–98. doi: 10.4049/jimmunol.2100969

[76] Korte R, Lepski S, Brockmeyer J. Comprehensive peptide marker identification for the detection of multiple nut allergens using a non-targeted LC-HRMS multi-method. Analytical and bioanalytical chemistry 2016;408:3059–69. doi: 10.1007/s00216-016-9384-4

[77] Skypala I, Bauer M, DunnGalvin A, Venter C. The Challenges of Managing Multiple Food Allergies and Consequent Food Aversions. The journal of allergy and clinical immunology In practice 2022;10:35–44. doi: 10.1016/j.jaip.2021.10.044

[78] Ellis JS, Guloglu FB, Zaghouani H. Presentation of high antigen-dose by splenic B220(lo) B cells fosters a feedback loop between T helper type 2 memory and antibody isotype switching. Immunology 2016;147:464–75. doi: 10.1111/imm.12579

[79] Martin GH, Gregoire S, Landau DA, Pilon C, Grinberg-Bleyer Y, Charlotte F, et al. In vivo activation of transferred regulatory T cells specific for third-party exogenous antigen controls GVH disease in mice. Eur J Immunol 2013;43:2263–72. doi: 10.1002/eji.201343449

[80] Karlsson MR, Kahu H, Hanson LA, Telemo E, Dahlgren UI. Tolerance and bystander suppression, with involvement of CD25-positive cells, is induced in rats receiving serum from ovalbumin-fed donors. Immunology 2000;100:326–33. doi: 10.1046/j.1365-2567.2000.00050.x

[81] Wang LJ, Mu SC, Lin MI, Sung TC, Chiang BL, Lin CH. Clinical Manifestations of Pediatric Food Allergy: a Contemporary Review. Clin Rev Allergy Immunol 2022;62:180–99. doi: 10.1007/s12016-021-08895-w

[82] Motta CM, Califano E, Scudiero R, Avallone B, Fogliano C, De Bonis S, et al. Effects of Cadmium Exposure on Gut Villi in Danio rerio. International journal of molecular sciences 2022;23. doi: 10.3390/ijms23041927

[83] Villanaccia V, Ceppab P, Tavanic E, Vindignid C, Volta U. Coeliac disease: The histology report. Digestive and liver disease : official journal of the Italian Society of Gastroenterology and the Italian Association for the Study of the Liver 2011;43:S385–S95

[84] Oberhuber G, Granditsch G, Vogelsang H. The histopathology of coeliac disease: time for a standardized report scheme for pathologists. Eur J Gastroenterol Hepatol 1999;11:1185–94

[85] Marsh MN. Intestinal Manifestations of Food Hypersensitivity. New York: Raven Press; 1995.

[86] Kangwan N, Kongkarnka S, Boonkerd N, Unban K, Shetty K, Khanongnuch C. Protective Effect of Probiotics Isolated from Traditional Fermented Tea Leaves (Miang) from Northern Thailand and Role of Synbiotics in Ameliorating Experimental Ulcerative Colitis in Mice. Nutrients 2022;14. doi: 10.3390/nu14010227

[87] Rocamora-Reverte L, Melzer FL, Wurzner R, Weinberger B. The Complex Role of Regulatory T Cells in Immunity and Aging. Frontiers in immunology 2020;11:616949. doi: 10.3389/fimmu.2020.616949

[88] El Ansari YS, Kanagaratham C, Burton OT, Santos JV, Hollister BA, Lewis OL, et al. Allergen-Specific IgA Antibodies Block IgE-Mediated Activation of Mast Cells and Basophils. Frontiers in immunology 2022;13:881655. doi: 10.3389/fimmu.2022.881655

